# HDAC inhibition via suberoylanilide hydroxamic acid (SAHA) ameliorates Doxorubicin-induced cardiotoxicity

**DOI:** 10.64898/2026.01.12.698976

**Authors:** Benay Eksi, Daniel Finke, Synje Michel, Jannek Brauer, Markus B Heckmann, Mohsen Valadan, Leonard M. Schanze, Vighnesh Sunder, Hugo A. Katus, Norbert Frey, Johannes Backs, Lorenz H. Lehmann

## Abstract

**Background:** Anthracycline-induced cardiotoxicity remains a major limitation of cancer therapy, and effective preventive strategies are lacking. Topoisomerase IIβ (Topo IIb) has been implicated as a central driver of this toxicity, suggesting that epigenetic regulators may interfere with pathological cardiac response.

**Methods and Results:** Here, we show that doxorubicin promotes Topo IIb accumulation at cardiomyocyte gene promoters (e.g., *Actc1, Myl2*, and *Myh7)* overlapping myocyte enhancer factor 2 (MEF2) binding sites and enhances MEF2-dependent transcription. This response is attenuated by the pan–histone deacetylase (HDAC) inhibitor suberoylanilide hydroxamic acid (SAHA). SAHA-mediated cardioprotection requires class IIa HDACs, as genetic loss of HDAC4 abolishes its effect. Mechanistically, SAHA induces acetylation of the chaperone 14-3-3, disrupting its interaction with HDAC4/5, promoting their nuclear accumulation, and repressing MEF2-driven transcription. In vivo, SAHA mitigates doxorubicin-induced cardiotoxicity.

**Conclusion:** These findings identify HDAC inhibition as a cardioprotective repurposing strategy and reveal a mechanistic link between epigenetic regulation and anthracycline-associated cardiotoxicity.

## Introduction

Despite decades of widespread clinical use, anthracyclines such as doxorubicin remain among the most frequently prescribed anticancer agents, however the molecular mechanisms underlying doxorubicin-induced cardiotoxicity (Doxo-Tox) are still incompletely understood. Current evidence implicates induction of reactive oxygen species (ROS) with subsequent mitochondrial dysfunction and increased apoptosis. This process is probably driven by poised topoisomerase IIb (Topo IIb) and impairments in DNA damage response^1^. To date, the iron chelator dexrazoxan is the only FDA-approved cardioprotective agent so far^2^. Thus, a deeper understanding of the molecular mechanisms is needed to improve safety of anthracycline-treated cancer patients.

Epigenetic modifiers, such as inhibitors of histone deacetylase activity (HDACi), have served as a novel class of anticancer drugs within the last years^3^. Beyond their antiproliferative potential^4^, increasing evidence is coming from preclinical models that HDACis may also exert cardioprotective effects in diverse models of cardiac stress^5,6^. Notably, the pan-HDAC inhibitor suberoylanilide hydroxamic acid (SAHA) has recently been explored as a potential adjunctive therapy to increase antiproliferative potential of classical treatment strategies of malignant diseases. SAHA has been incorporated into combination therapies to treat non-hodgkin lymphoma^7^ or multiple myeloma^8^ or sarcoma^9^. Elucidation of a potential cardioprotective mechanism of SAHA may therefore promote such co-treatment strategies to eventually mitigating cardiotoxic effects of anthracyclines.

HDACs are classified in multiple subclasses, among which class I and class II HDACs are supposed to be strongly involved in cardiac hypertrophy and pathological cardiac remodeling^10^. Class I HDACs are well established drivers of cardiac pathological remodeling^11,12^. Giving their high deacetylase activity of class I HDACs compared to class II HDACs, it is likely that gene expression of antihypertrophic genes is modulated via a direct inhibition of class I HDACs by HDACis^12^. However, class II HDACs are known as potent inhibitors of pathological cardiac remodeling. For instance, genetic deletion of the class II HDAC HDAC5 leads to a higher sensitivity to prohypertrophic stimuli in a genetic model of cardiac hypertrophy^13^. HDAC6 was shown to mediate the cardiac stress response upon Angiotensin II administration^14^. In contrast to class I HDACs (1, 2, 3 and 8), class II HDACs (HDAC 4, 5, 7 and 9) have posess minimal intrinsic deacetylase activity. Therefore, they are unlikely to represent direct enzymatic targets of HDACis^15^. Crystal structure analyses have shown that the HDACi SAHA binds selectively to the catalytic deacetylase pocket of the HDACs^16^, further supporting the notion that class II HDACs are regulated primarily through non-enzymatic mechanisms. Instead, they act signaling-responsive by shuttling from the nucleus into the cytosol^17,18^. The shuttling depends on their binding to the chaperon protein 14-3-3^19^. When retained in the nucleus, HDAC4 and 5 inhibit the transcription factor Myocyte Enhancer Factor-2 (MEF2)^20^, essential for cardiomyocyte hypertrophy and pathological cardiac remodeling. A redundant role for class II HDACs is shown for HDAC 5 and 9^13^, whereas HDAC9 is only poorly expressed in cardiomyocytes. A possible role of class II HDACs in translating antihypertrophic effects of HDACis remains unclear.

Here, we identify a previously unrecognized mechanistic link between Topo IIb-dependent DNA damage and MEF2-driven transcriptional remodeling in cardiomyocytes. We show that doxorubicin induces Topo IIb enrichment at MEF2-DNA binding sites, including promoters of cardiomyocyte-specific and remodeling-associated genes. We further demonstrate that SAHA suppresses MEF2 activity through a mechanism that requires HDAC4 and involves acetylation-dependent disruption of 14-3-3–class II HDAC interactions. Using pharmacological inhibition, genetic loss-of-function models, and in vivo cardiotoxicity assays, we establish that HDAC4 is essential for the cardioprotective effects of SAHA during anthracycline treatment. Together, our findings define the class II HDAC-14-3-3-MEF2 axis as a critical regulator of anthracycline-induced cardiac remodeling and reveal a mechanistic basis for HDAC inhibitor-based cardioprotective adjunctive therapy.

## Methods

### Animal experiments

All animal experiments were reviewed and approved by the Animal Experiment review board of Baden-Württemberg, Germany. Wildtype animals were purchased from Janvier. Genetically modified animals were bred in the central animal housing of Heidelberg University (IBF). C57BL6J mice at the age of 10 weeks were injected 8 times within 2 weeks with doxorubicin in a concentration of 3 mg/kg BW ± 12.5 mg/kg BW SAHA per injection. Cardiac function was assessed via echocardiography under isoflurane anesthesia (2% v/v). 10 weeks after the last injection, the animals were sacrificed and downstream analysis was performed. TMP195 was administered in 10 weeks-old WT animals (C57BL6J) in a dose of 15mg/kg BW per injection with total injections of 7 (Doxo, 3 mg/kg BW/per injection) within 2 weeks. Echocardiography under anesthesia were performed at three timepoints; at baseline before starting the treatment, right after the treatment phase and at the end of the experiment (9 weeks after the last treatment). Cardiomyocyte-specific HDAC4-deficient mice (HDAC4 cKO; C57BL6N) were obtained by crossing a floxed allele of HDAC4 in an a-MHC MerCreMer mouse line as described earlier^21^. WT littermates carried either a floxed allele and no MerCreMer (cre-) or no floxed allele but MerCreMer (cre+). The knockout was induced by Tamoxifen application. Tamoxifen was given intraperitoneally over a 10-day period, comprising 5 days of consecutive administration, a 1-day pause and 4 additional days of treatment. Each animal received a dose of 1 mg/day (10 µg/µl). The cre status of the control littermates is marked in the respective figures. 10-13 weeks old animals (female and male) received Doxo in a concentration of 3 mg/kg BW ± 12.5 mg/kg BW SAHA per injection with a total of 7 injections within 2 weeks. Echocardiography under anesthesia (2% isoflurane) was performed at baseline, right after the treatment phase and at the end of the experiment. The HDAC4-cKO animals were sacrificed 4 weeks after the last injection.

Animals were euthanized by cervical dislocation and organs were harvested and weight was assessed using a precision balance (Sartorius). Tibias were separated from the surrounding tissue by H_2_O_2_ digestion.

### Genotyping PCR

Genomic DNA for genotyping was isolated from ear punches using lysis buffer (NaOH, 0.5M EDTA, incubation for 20 min at 95 °C) and neutralization buffer (Tris-HCl, shortly mixed and centrifuged). REDextract PCR Ready Mix (Sigma-Aldrich) was used for the PCR setup. The following primer were used:

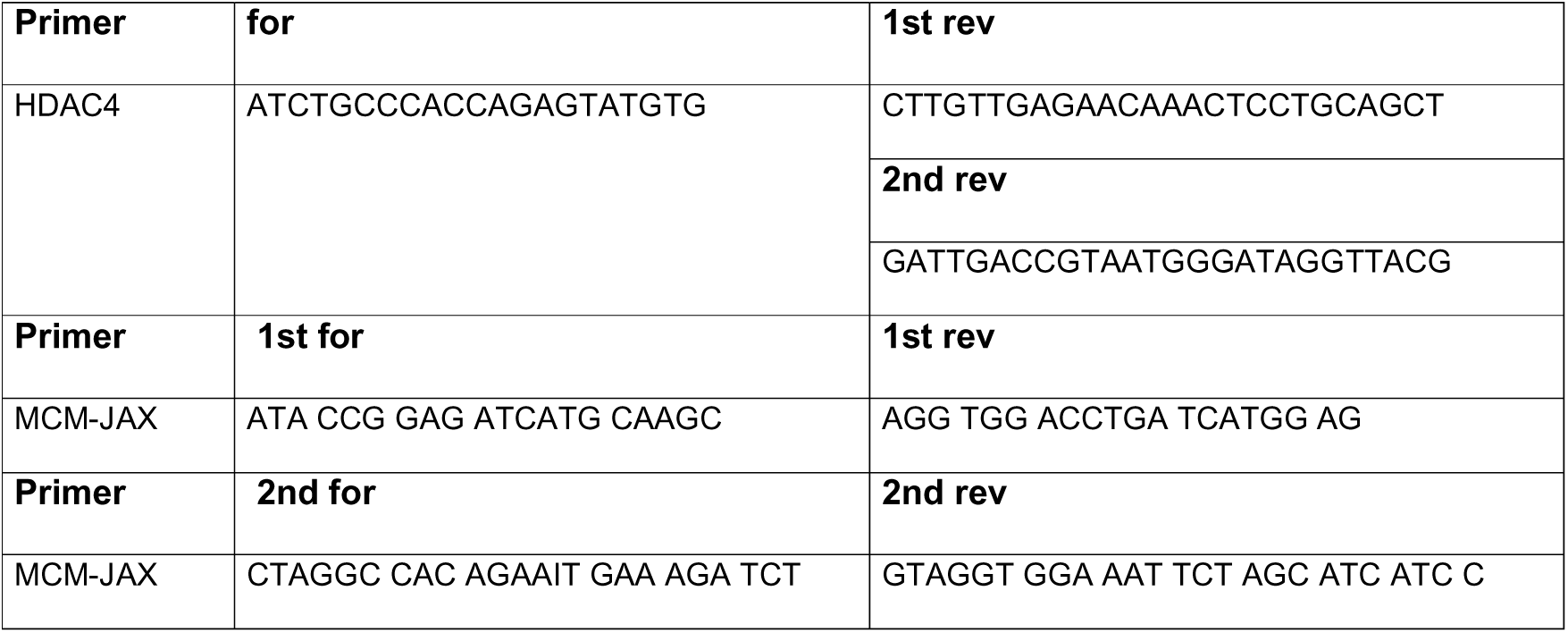

### Transthoracic echocardiography

Echocardiography was performed using a Visual Sonics vevo 2100 using a MX550D transducer on anesthetized animals (2% isoflurane). Prior to the echocardiography, the mice were shaved (asid med shaving creme). Measurements of left ventricular ejection fraction were done in M-Mode images from the parasternal short axis at the papillary muscle level using Vevo Lab software (version 5.7.1). At least four consecutive beats were used for the measurements.

### Histology

Hearts were fixed in 10% paraformaldehyde and dehydrated in ethanol. Afterwards they were embedded in paraffin. Sections were stained with sirius red. Staining was performed on 1-μm slices as explained before^22^.

In brief, cuts were deparaffinized, rehydrated and incubated with hematoxylin (5min, Morphisto) and eosin (2min, Leica), or sirious red (Morphisto) respectively.

Images were taken with the Zeiss Axio slide scanner Axio Scan.Z1 (Carl Zeiss, Germany) in brightfield mode with a 4.4 µm/pixel sampling (tile size 1600 × 1200 pixels; field of view 7.04 × 5.28 mm) and 24-bit RGB color. Images were acquired using 1 × 1 binning. No additional image processing was applied prior to quantification, apart from insertion of scale bars and global adjustments of brightness and contrast for figure presentation.

#### Cell culture and transfection assays

COS cells were maintained in DMEM with FBS (10%), L-glutamine (2 mM) and penicillin-streptomycin. Transfection of COS cells was performed with Lipofectamine 2000 (ThermoFisher) according to the manufacturer’s instructions.

#### Western Blot Analysis

Proteins from cultured cardiomyocytes or COS cells were isolated and Western blot analysis was performed according to the protocols described before^23^. Equal amounts of protein extracts were separated with SDS-PAGE and transferred to a PVDF membrane (Millipore). The utilized primary antibodies were anti-FLAG-tag (Millipore), anti-myc (CellSignaling), anti-14-3-3 (abcam), anti-HDAC4 (abcam, ab12172) and GAPDH (CellSignaling, 2118S). Primary capture antibody incubation was followed by secondary detection anti-mouse and anti-rabbit antibody exposure. The detection antibodies were labeled with horseradish peroxidase (HRP) and quantified by subsequent ECL detection.

#### Adenovirus production

Adenovirus harboring myosin enhancer factor 2 (MEF2) luciferase was purchased from Seven Hills Bioreagents. Adenovirus harboring HDAC4/HDAC5 constructs were described previously^24^. After generation, the adenoviruses were amplified, purified with the Adeno-X Purification Kit (BD) and their infectious units per µl were determined with the Adeno-X Rapid Titer Kit (BD).

#### Culture of neonatal rat ventricular cardiomyocytes (NRVMs)

NRVMs were isolated from 1– to 2-day-old Sprague Dawley rats as previously described^24^. After isolation, NRVMs were cultivated in complete medium with 10% FBS, 2 mM L-glutamine and 1% penicillin-streptomycin. NRVMs were infected with ad-HDAC4 virus 24h after plating. Cells were grown for additional 24h in fresh complete medium. The medium was then replaced with serum-free medium 24h prior to adding test drugs as indicated in the experiments. Cells were then fixed for immunocytochemical staining. For MEF2 luciferase reporter assays, adenoviruses harboring MEF2 luciferase were applied to cultured cardiomyocytes for 24h before treating with indicated chemicals. For overexpression of HDAC4, the HDAC4-FLAG adenovirus was added after cultivating NRVMs for 24 h. After washing, the cells were treated with the indicated chemicals for 4 h.

#### Luciferase Assay

Treated NRVMs (Doxo 1µM, 4h and SAHA 3µM, 6h) were collected in 150µl lysis buffer (Promega) and transferred to a 96-well plate. Luciferase and Renilla was each added to the samples in a 1:1 dilution. Luciferase signal reads were measured as end-points by the Microplate Reader (*Perkin Elmer*).

#### Chromatin Immunoprecipitation

NRVMs (10 million cells) were treated with doxorubicin (1 µM) for 3 h and crosslinked with formaldehyde (1%, diluted in PBS) for 10 min. The reaction was quenched by administration of glycine to a final concentration of 250mM. Chromatin was isolated and sheared using Bioruptor (Diagenode) using 3×10 cycles (High Power, 30 sec power – 30 sec pause) as descried earlier.^25^

The immunoprecipitation was done with the iDeal ChIP-seq Kit for Histones (Diagenode) using IP-Star (Diagenode) following the manufacture’s guidelines. Topoisomerase IIb antibody (Santa Cruz, A-12, sc-365071, 3.5µg) was used for the chromatin pulldown (3,0µg chromatin).

### ChIP-seq library preparation and next-generation sequencing

ChIP-seq libraries have been prepared using the NEBNext® Ultra™ II DNA Library Prep Kit following the manufacturer’s instructions (New England Biolabs). Bioanalyzer (Agilent Genomics) analysis confirmed the libraries’ quality. Four samples have been sequenced on one lane as 100 bp single-end reads using Illumina HiSeq 2000 (Illumina, San Diego, USA).

#### Transcriptional Analysis

##### RNA extraction and quantitative PCR

Total RNA was isolated from cells or homogenized cardiac tissue using Qiazol (Qiagen). Samples were incubated with chloroform at room temperature and centrifugated for 15 min at 14.000 rpm. The aqueous phase was mixed with isopropanol and glycogene (ThermoFisher) and incubated at –20°C for at least 1h. The solution was centrifugated for 30 min in order to precipitate the total-RNA. Pellets were washed twice with 70 %v/v ethanol. RNA concentration was measured by photometry (Nanodrop, ThermoFisher) and purity was verified (OD_260_/OD_280_□>□1.8). Complementary DNA (cDNA) synthesis of 1µg of RNA was carried out using a cDNA synthesis kit (Biozym).

Sybr Green reagents (ThermoFisher) were used for quantitative PCR using the standard curve settings. The following primer used for the detection of gene expressions:

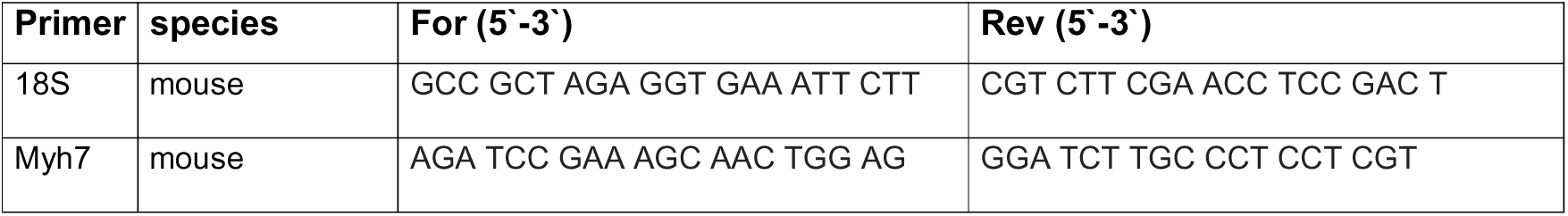

#### Bulk-RNA-seq

Total RNA was isolated from cardiac tissue using Qiazol (Qiagen). mRNA-library preparation and RNA-seq was performed by BGI genomics (Hong Kong) on a DNBDEQ platform (20 million reads, 100bp, paired end). 3 animals/ group were analyzed in WT Doxo and SAHA treatments (Figure 3). 5 animals/ group were analyzed in RNA-seq of HDAC4-cKOs and control littermates (Figure 5).

#### Immunofluorescence – HDAC4 Localization

NRVMs were fixed in 4 % paraformaldehyde (Sigma-Aldrich) in PBS, permeabilized with 0.3 % Triton X-100 (Sigma-Aldrich) for 10 min at room temperature and blocked for 60min with 5 % goat serum (PAA) in PBS. Primary antibodies for α-actinin (Sigma) and FLAG were applied in PBS containing 5% goat serum or BSA and 0.05% Triton X-100 for 1h. Cells were treated with the secondary antibody (Texas Red-coupled anti-mouse-antibody and Alexa fluor 488 coupled anti-rabbit-antibody) in PBS containing 5 % goat serum and 0.05% Triton X-100 for 1 h. Nuclei were stained with DAPI (Invitrogen). For quantification blinded >10 fields of view were recorded and the percentage of cells containing HDAC4 in the cytosol for each taken picture were counted.

#### Mammalian 2-hybrid assay

A mammalian expression vector encoding the GAL4 DNA-binding domain fused to the amino-terminus of human HDAC4 (amino acids 2–740) was generated in the pM expression vector (Clontech). COS cells were transiently transfected with vectors for GAL4-HDAC4, VP16–14-3-3 and a luciferase reporter gene under the control of 5 copies of a GAL4 DNA-binding site (5 × UAS-luciferase). The expression vectors were transfected in the absence or presence of a construct encoding protein kinase D (PKD). Transfection efficiency was controlled by co-transfection of Renilla luciferase. 24 h after transfection, cells were harvested and luciferase levels were determined as described above.

#### Coimmunoprecipitation and immunoblotting

COS cells were harvested 24 h after transfection in Tris (50 mM, pH 7.4), NaCl (150–900 mM), EDTA (1 mM) and Triton X-100 (1 %) or in RIPA buffer supplemented with protease inhibitors (Complete; Roche Diagnostics) and PMSF (1 mM). Cells were further disrupted by passage through a 25-gauge needle and cell debris was removed by centrifugation.

FLAG-tagged proteins were immunoprecipitated with M2-agarose conjugate (Sigma-Aldrich) and thoroughly washed with lysis buffer.

NRVMs were treated with SAHA (3µM for 1h) for immunoprecipitation of acetylated proteins. Pan-acetyl antibody bound beads (Immune Chem, ICP0388) were used using the manufacture’s guidelines with minor adjustments. Briefly, 40µl of the anti-acetyl lysine agarose was washed three times in 1ml TBST, twice in 1ml of 0.1M NaH_2_PO_4_/ 1M NaCl, followed by one wash in 1ml TBST. 1.000 mg of protein was incubated with the washed pan-acetyl lysine beads overnight at 4°C. After incubation, the protein-beads samples were washed four times with TBST.

Bound proteins were resolved by SDS-PAGE, transferred to PVDF membranes and immunoblotted as indicated by a monoclonal anti-FLAG antibody (M2; Sigma-Aldrich) in overexpression experiments. A 14-3-3 antibody (abcam, rabbit, ab125032) was used for immunoblotting after pan-acetyl pulldown. Proteins were visualized with a chemiluminescence system (Santa Cruz Biotechnology Inc.).

#### siRNA treatment of NRVMs

Reverse transfection of siRNA duplexes into NRVMs using Lipofectamine 2000 transfection reagent (ThermoFisher Scientific, Waltham, MA) has been described before^26^. NRVMs (1.5 × 10^6^ cells) were briefly suspended in medium containing 10% FCS, incubated with 100 nM siRNA and transfection reagent, followed by plating in 12-well plates overnight. Medium was changed the next day. The transfection was repeated twice. The sequence of siRNA targeting rat HDAC1-11 are shown in the table.

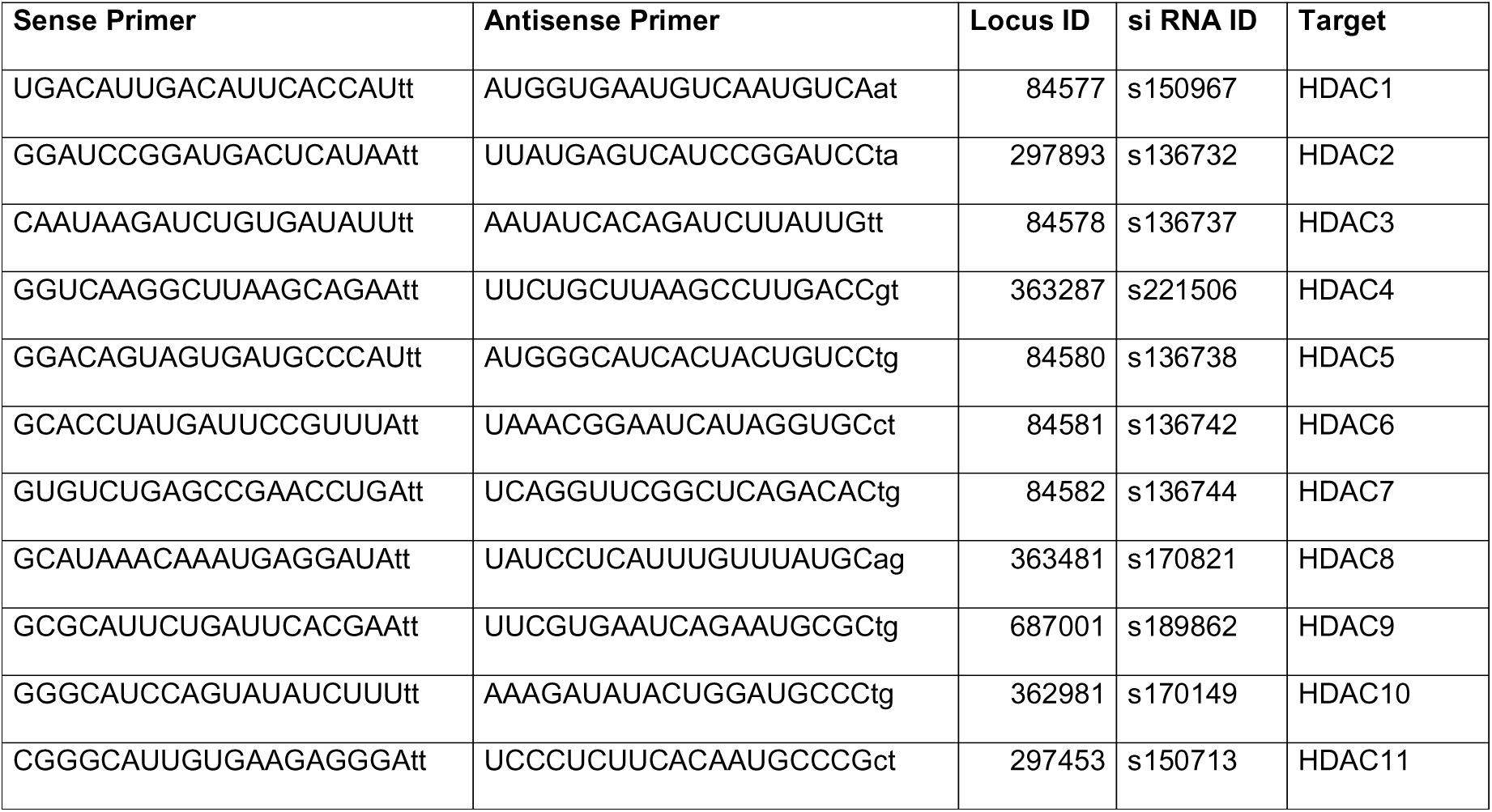

A non-targeting sequence (cat#12935300, Thermo Fisher) was used as a control siRNA.

### Used Primer for realtime PCR

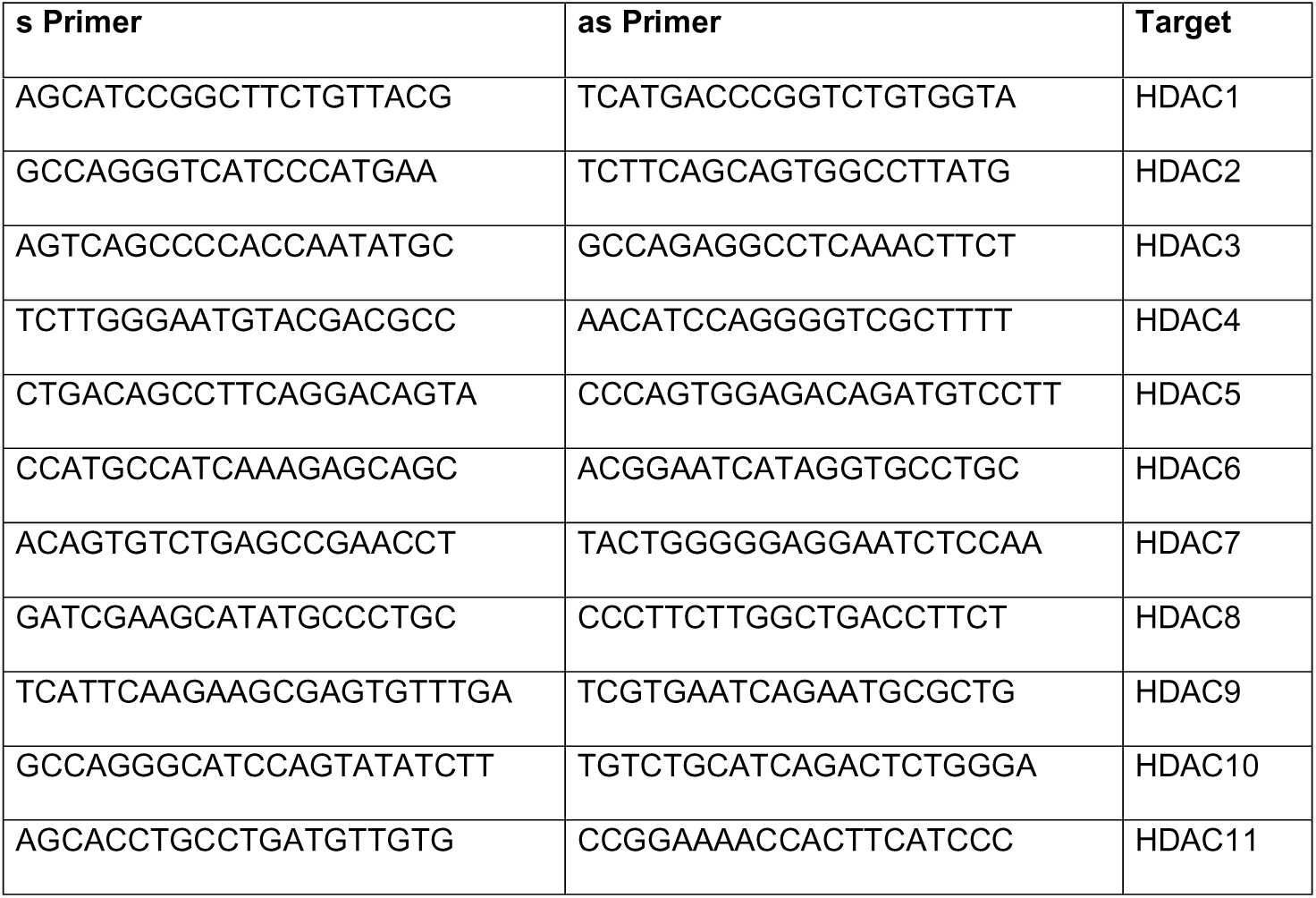

### Statistical Analyses

For all statistical analyses, the software GraphPad Prism (version 10.4.2) was used. Comparison between two groups was performed using the Mann-Whitney-U-test while for comparison of more than two groups, one way-ANOVA with Bonferroni correction was used to assess statistical differences. A p-value less than 0.05 was considered statistically significant. Number of replicates per group are stated in the figure legends or the corresponding text.

### Analysis of histology

Analysis of histological images for fibrosis measurements were performed using ImageJ (version 1.54). Images were converted in 8-bit black and white images. The same intensity threshold to assess fibrosis was used in all analyzed images. One heart section per mouse was stained and 5 Regions of Interests (ROI) per heart section were selected randomly. For each ROI, the area of fibrosis was measured (same cutoff for all) and normalized to the whole area of the ROI (in %). The number of replicates analyzed per group are shown in the figure legend. Each data point in the figures represents a single ROI. For statistical comparison one-way ANOVA with Bonferroni correction was performed using GraphPad Prism version 10.4.2.

### Analysis of ChIP-seq

FASTQ data were annotated to the rat genome (rn5) using bowtie2 (version 2.3.4.1). Only uniquely mappable reads have been subjected to further analysis. Peak calling was performed with MACS2 (version 2.1.1) with a selected p-value of 0.0001 and normalization to the respective input file. Differentially enriched peaks were calculated with DESeq2 (version 1.24.0). An FDR < 0.1 was used as a cutoff.

Heatmap and profiles of genomic enrichment were illustrated in Deeptools (version 3.2.1). Exemplary sequencing tracks were visualized in the IGV browser (version 2.13.1) as reads per kilobase per million (RPKM) and 50-bp binning. Files were created from BAM files via the bamCoverage tool. The input file was subtracted from ChIP and files were averaged (bigwigAverage). Principal component analysis (PCA) was performed with multiBamSummary (version 3.5.4), with a binning size of 10.000bps and distance between bins of 0 (**Suppl. Figure 1A**). Annotated peak numbers and fraction of reads in peaks (FRiP) scores are listed in **Suppl. Table 1**. Scatterplots and pie charts were created in GraphPad prism (version 10.4.1). ChIP-seq peaks were annotated to genomic locations and features (e.g., transcription start site (TSS), intergenic, intragenic) with HOMER (v. 4.10) using the annotatePeaks.pl command.

### Analysis of RNA-seq

FASTQ files of RNA-seq reads were aligned to the mouse genome version mm9. Counts per transcript were extracted with featureCounts (Rsubread). Reads were normalized to reads per kilobase per million (RPKM) within the DESeq2 package (version 1.40.2).

PCA of normalized RNA-seq expression profiles was conducted prior to downstream analyses to assess treatment-related clustering and sample quality within the Deseq2 package. Differentially expressed genes (DEG) were calculated between two groups (WT Doxo vs NaCl) using the DESeq2 algorithm. The set of DEGs (FDR < 0.05) was visualized across all treatment groups in a heatmap using the Chi-squared (Chisq) distance on expression profiles for clustering in heatmap3 package (version 1.42.1).

### Analysis of external data

Data from hiPSC-cells were obtained from (GEO ID <u>GSE106297</u>)^27^ and re-analyzed with normalized count matrices (FPKM). Reads were aligned to the human genome (hg38) for gene annotation. Statistical analysis was performed by one-way ANOVA with Bonferroni correction using GraphPad Prism (version 10.4.2).

### Data availability

Raw and annotated ChIP-seq Topo IIb data have been submitted to the EMBL-EBI database (accession number: E-MTAB-16326). MEF2 ChIP-seq was published earlier (accession number: E-MTAB-6213). RNA-seq data are deposited under the accession number: E-MTAB-16361.

## Results

### Genomic topoisomerase IIb binding identifies cardiac doxorubicin target genes

Doxorubicin is affecting topoisomerase IIb (Topo IIb) activity. To identify downstream targets of doxorubicin, Topo IIb ChIP-seq was performed in neonatal rat ventricular cardiomyocytes (NRVM), that were treated with doxorubicin. We were able to identify genomic regions significantly gaining Topo IIb enrichment after doxorubicin treatment (n = 211 peaks, FDR < 0.1) indicative of a specific response on double strand breaks. These regions comprise cardiomyocyte-specific promoters, e.g., of *Myh7*, *Myl2* and *Actc1* (**Figure 1A-E, Suppl. Figure 1B-D**) and share a high enrichment of the transcription factor myocyte enhancer factor 2 (MEF2), known to be a major mediator of pathological cardiac remodeling (**Figure 1A, E**). From 1148 called peaks in Doxo treated NRVMS (at least on replicate), we found 1012 (88%) peaks overlapping with at least one MEF2-peak. A total of 180 peaks were recalled in all replicates of Topo IIb– and MEF2-ChIP-seq as common targets (**Figure 1F**).

**Figure 1.**
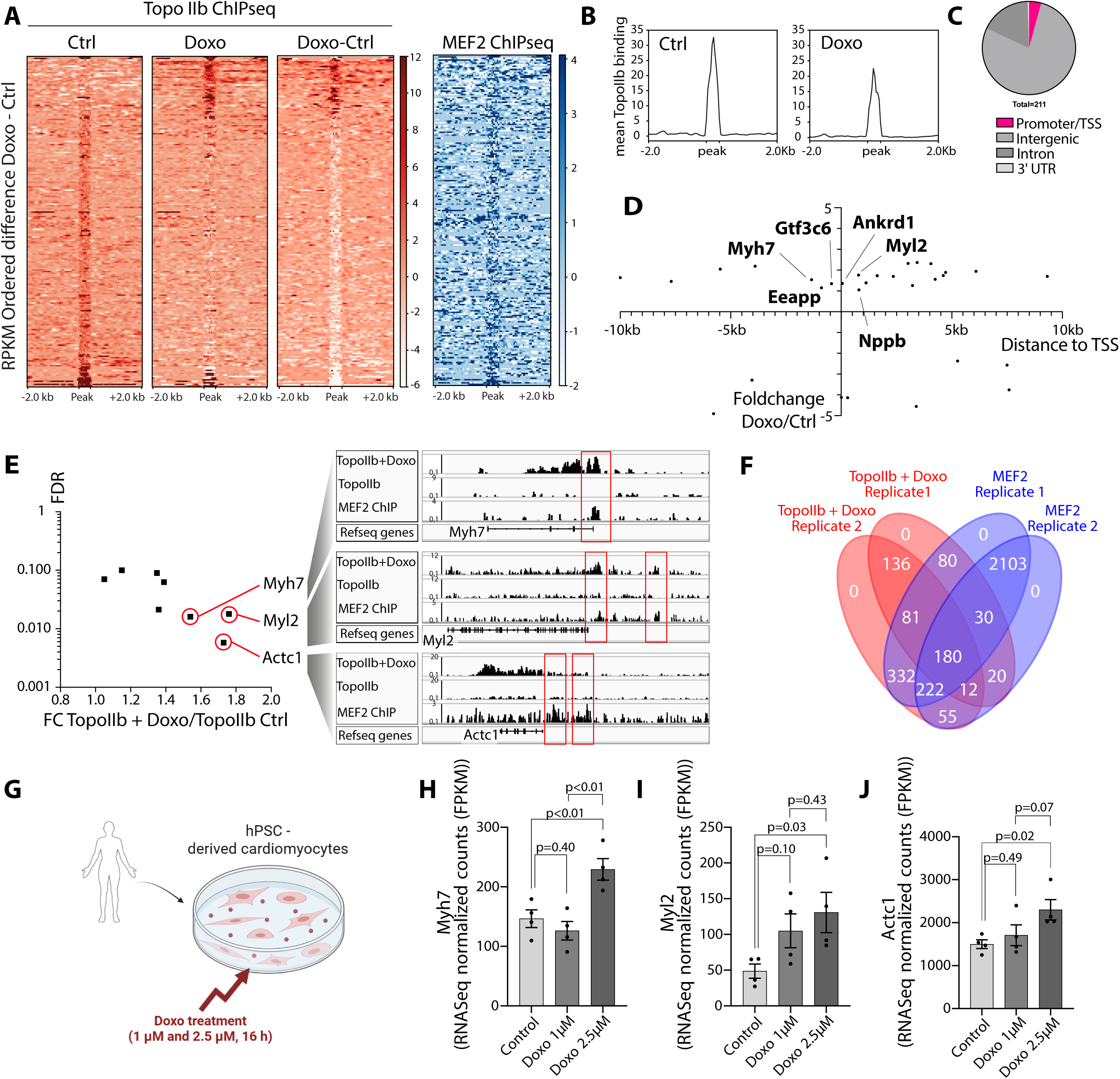
(**A**) Heatmaps of topoisomerase IIb (Topo IIb) ChIP-seq enrichment in control (Ctrl) and doxorubicin-treated (Doxo) NRVMs at differentially enriched peaks (Doxo vs. Ctrl, two averaged replicates/treatment, FDR<0.1). Rows correspond to genomic regions. These regions were plotted in the same order for Ctrl (first panel), doxorubicin-treated cells (second panel) and for the difference Ctrl subtracted from Doxo (Doxo-Ctrl) in the third panel. The rightmost heatmap shows MEF2 ChIP-seq enrichment over the same ordered set of regions. **(B)** Average Topo IIb ChIP-seq enrichment over all differentially enriched peaks in Ctrl and Doxo-treated NRVMs (±2 kb around the peak). **(C)** Distribution of Doxo-induced Topo IIb peaks to genomic locations. The pie chart shows the respective fraction of peaks located in promoters/transcriptional start sites (TSS), intergenic regions, introns or 3′ UTRs. **(D)** Scatterplot highlighting the differential Topo IIb bound regions after Doxo treatment with the highest foldchange (FC) Doxo/Ctrl (y-axis). The x-axis indicates the genomic distance of the peak to the TSS (in kb). **(E)** Scatterblot of the most regulated Doxo-induced Topo IIb peaks according to FC and false discovery rate (FDR). Exemplary ChIP-seq tracks are shown for Topo IIb (Ctrl and Doxo) and MEF2 for promoter regions of *Myh7*, *Myl2* and *Actc1.* **(F)** Topo IIb was binding after Doxo-treatment in NRVM on specific sites (409 induced binding-sites in the two replicates). A high fraction (180) was overlapping with MEF2-binding sites (all replicates). **(G)** Experimental setting of treating human pluripotent stem cells (hPSC) derived cardiomyocytes (Maillet et al., 2016). Re-analysis of published RNA-seq data in hPSC-derived cardiomyocytes treated with doxorubicin (vehicle, 1 µM, 2.5 µM; 16 h). Gene expression for *Myh7* **(H)**, *Myl2* **(I)** and *Actc1* **(J)** reveals a significant increase after doxorubicin treatment (2.5µM). Normalized values (FPKM) for *Myh7, Actc1* and *Myl2*; points are replicates (2 sets of duplicates, n=4) and bars show mean ± SEM.

### Target genes are regulated in doxorubicin-treated hiPSC

To verify transcriptional dysregulation of the identified genes in human samples, we re-analyzed the publicly available RNA-seq dataset of hiPSC-derived cardiomyocytes exposed for 16 h to vehicle, 1 µM or 2.5 µM doxorubicin (2 sets of duplicate samples per condition) (**Figure 1G**). We visualized *Myh7*, *Actc1* and *Myl2* by plotting normalized counts (FPKM) across all three conditions (**Figure 1H-J**). All three genes showed treatment-associated changes relative to vehicle in a dose-dependent manner. *Myh7* was significantly upregulated at a dose of 2.5 µM compared to the control and 1 µM doxorubicin (p=0.002), while 1 µM of doxorubicin did not lead to changes compared to the control (p=0.401). Genes, such as *Actc1* and *Myl2* showed similar effects displaying significant changes at a dose of 2.5 µM compared to the control. We therefore conclude that these genes could serve as potential indicators for anthracycline-dependent Topo IIb-guided MEF2 activation.

#### SAHA inhibits MEF2 activity in cardiomyocytes

Since the HDACi SAHA is known to protect from pathological cardiac remodeling associated with pressure overload, we assessed its potential for re-purposing. Interestingly, in luciferase reporter assays, we found a concentration-dependent inhibition of MEF2 by SAHA treatment (**Figure 2A**). The activity of MEF2 in turn was increased by doxorubicin application and could be inhibited by SAHA (**Figure 2B**). Given the fact that SAHA is a pan HDAC inhibitor interacting with the deacetylase domain, we evaluated MEF2 activity after single knockdown of HDAC1-11 using siRNA to identify the SAHA target HDAC and the strongest MEF2 inhibiting HDAC (**Figure 2C**). Efficient knockdown of selected HDACs was confirmed by real-time PCR, using HDAC specific primer, respectively (**Suppl. Figure 2**). Intriguingly, the effects on MEF2 diverge markedly between different HDACs. HDAC3+6 knockdown reduced MEF2 activity, whereas the knockdown of the class II HDACs HDAC4+5 were found to induce the activity. Combinational knockdown of HDAC4+5 showed a synergistic effect with an exponentially induction of MEF2 activity. Vice versa, HDAC3+6 did not show additional synergistic effects (**Figure 2D**). Knockdowns of HDAC4+5 were able to mitigate SAHA-driven MEF2 inhibition, indicating that SAHA-driven MEF2 repression is at least partially dependent on class II HDACs 4+5. In contrast, HDAC3+6 knockdowns did not additionally repress MEF2 in the presence of SAHA, indicating that HDAC3+6 are potential direct targets of SAHA. Immunohistology of FLAG-tagged HDAC4 showed nuclear export of HDAC4 after doxorubicin, that is attenuated by administration of SAHA (**Figure 2E-F**). Nuclear localization of class II HDACs is known to mediate cardioprotective effects, including MEF2 inhibition.

**Figure 2.**
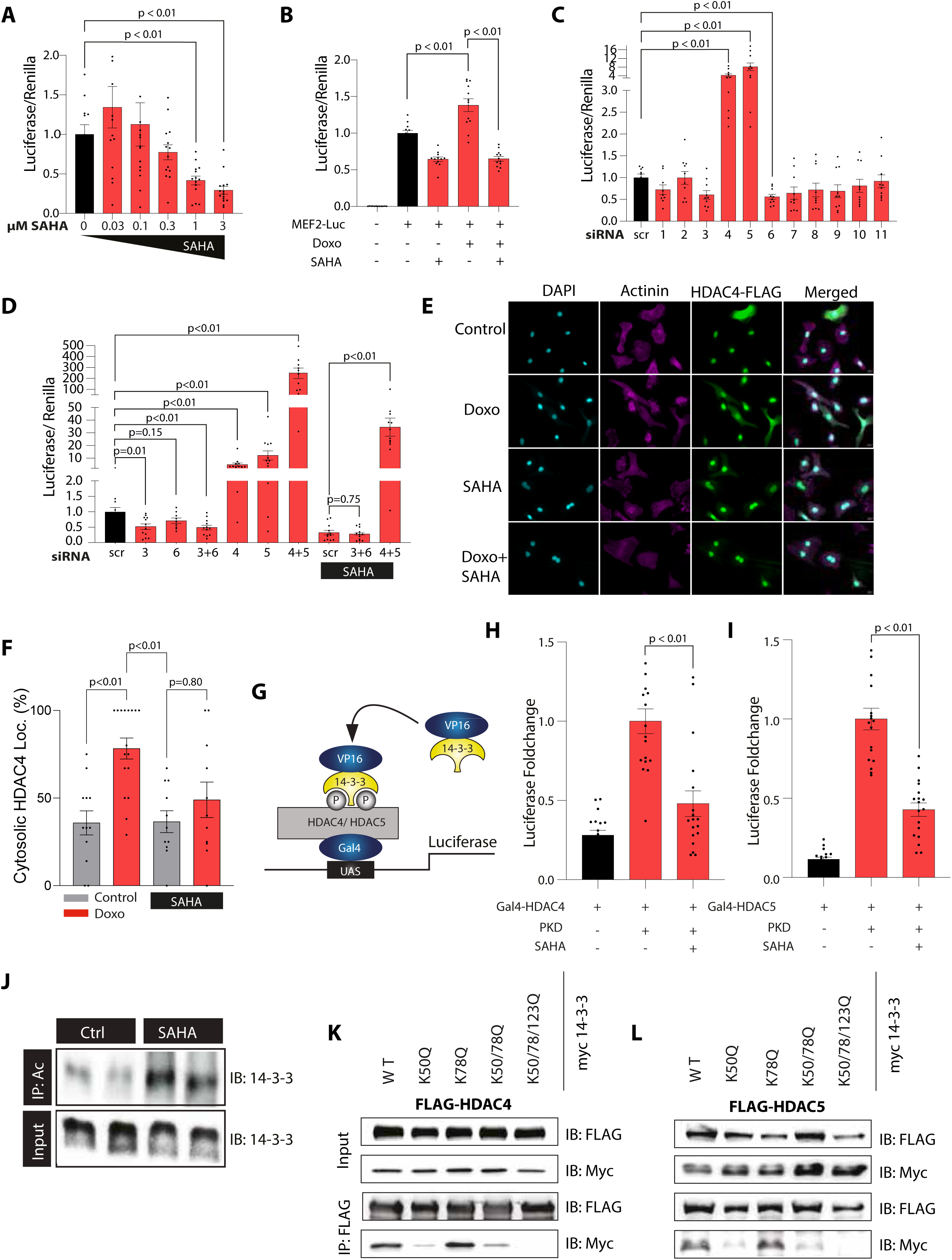
(**A**) Dose response of suberoylanilide hydroxamic acid (SAHA) on myocyte enhancer factor 2 (MEF2) activity measured via Luciferase Assays. The assays are normalized to Renilla luminescence (Luciferase/Renilla). P-values were calculated via Anova with Bonferroni posttest. Bars show mean ± SEM of 14-17 measurements per group from 3 independent experiments **(B)** Doxorubicin (300 nM) increases MEF2 activity, which is attenuated by co-treatment with SAHA (3 µM), Anova with Bonferroni posttest. Bars show Luciferase/Renilla values (mean ± SEM) of 12 measurements per group from 3 independent experiments. **(C)** siRNA driven knockdown of HDAC1-11 in NRVMs as indicated. MEF2 activity is shown as Luciferase/Renilla. 9-10 measurements/ group from 2 independent experiments, Anova with Bonferroni posttest. **(D)** Knockdown of HDACs as indicated and treatment with SAHA (3 µM). HDAC3/6 as well as HDAC4/5 were knocked down together, compared to the single knockdowns, respectively. 9-12 measurements/ group from 3 independent experiments, Anova with Bonferroni posttest. **(E)** Representative immunofluorescence images to detect cytosolic and nuclear HDAC4 localizations of NRVMs treated with doxorubicin (300mM) ± SAHA (3 µM) stained with DAPI (cyan), α-actinin (magenta) and HDAC4 (green). **(F)** Quantification of the relative cytosolic HDAC4 localization in percent. Each data point represents the percentage of cells with cytosolic HDAC4 localization within one image (Control, n=12; Doxo, n=16; SAHA, n=11; Doxo+SAHA, n=11). Anova with Bonferroni posttest. **(G)** Scheme of the used luciferase reporter assay to evaluate the binding of class II HDACs and 14-3-3 after treatment with SAHA. VP16-14-3-3 was overexpressed together with HDAC4/5-Gal4 and a UAS-reporter. **(H)** Relative Luciferase Activity of HDAC4-14-3-3 binding stimulated via PKD co-expression and inhibited with SAHA (3 µM). 18 measurements from 3 independent experiments. Mann-Whitney U-test. **(I)** Relative Luciferase Activity of HDAC5-14-3-3 binding stimulated via PKD co-expression and inhibited with SAHA (3 µM). 18 measurements from 3 independent experiments. Mann-Whitney U-test. **(J)** Immunoprecipitation of endogenous pan-acetylation sites ± SAHA pretreatment (3 µM) in NRVMs as indicated. Immunoblotting in pulldown and input for 14-3-3 is shown. IP: Immunoprecipitated antibody; IB: Immunoblotted antibody. **(K)** Immunoprecipitation of FLAG-tagged HDAC4 and **(L)** Immunoprecipitation of FLAG-tagged HDAC5 with co-expression of myc-tagged 14-3-3 chaperon with the indicated targeted mutations. IP: Immunoprecipitated antibody; IB: Immunoblotted antibody.

#### SAHA leads to 14-3-3 hyperacetylation and inhibits 14-3-3 – class II HDAC binding

The binding between the chaperon 14-3-3 and class II HDACs is a crucial factor for the cytosolic export of class II HDACs^19^. Using the fusion proteins 14-3-3-VP16 and HDAC4/5-Gal4 in control of the Upstream Activation Sequence (UAS-Enhancer) in a mammalian two hybrid assay, we validated SAHA-dependent disruption of the 14-3-3-HDAC binding (**Figure 2G-I**). Since SAHA may modulate chaperone 14-3-3 activity and class II HDAC, we analysed a SAHA-dependent modification of 14-3-3.

Therefore, we studied whether endogenous 14-3-3 may be acetylated after treatment with SAHA as a non-histone target. By immunoprecipitation for pan-lysine residue acetylation and immunoblotting for endogenous 14-3-3, we found a profound increase in acetylation of 14-3-3 in NRVMs if pretreated with SAHA (**Figure 2J**). In order to mimic acetylation of 14-3-3, we mutated the specific lysine residues which were earlier suggested to be hyperacetylated upon SAHA treatment^28^. Doing so, we found a profound reduction of the binding between 14-3-3 and HDAC4 (**Figure 2K**) or HDAC5 (**Figure 2L**) in co-immunoprecipitation experiments. These data indicate that mainly K50 is responsible for acetylation effects. In contrast, the lysin residue 78 alone could hardly show an effect on the protein-binding in cardiomyocytes.

#### SAHA attenuates doxorubicin-induced cardiotoxicity

To assess the in-vivo-relevance of the cardioprotective effects of SAHA, C57BL6J mice, aged 10 weeks, were co-treated with doxorubicin (total of 24 mg/kg in 2 weeks) and SAHA (100 mg/kg in 2 weeks). The mice were monitored for up to 10 weeks to investigate the long-term effects on pathological gene expression and cardiac function in anesthetized mice (Isoflurane 2%) using echocardiography **(Figure 3A).** The treatments had no effect on the survival rate of the animals **(Figure 3B).** Doxorubicin-treated mice revealed a moderate decrease in heart weight normalized to tibia length **(Figure 3C)** and a significant reduction in bodyweight **(Figure 3D, Suppl. Figure 4A)**. Representative images of echocardiography are depicted in **Figure 3E**. The left ventricular ejection fraction (LVEF) and heart rate demonstrated a mild but consistent decline in doxorubicin-treated mice compared to the control group **(Figure 3E-G).** The combination therapy of doxorubicin and SAHA preserved cardiac function in the treated mice, as indicated by less pronounced changes in LVEF relative to the doxorubicin-only group **(Figure 3F)**. Cardiac diameters (mainly LVEDD) showed slightly smaller values in the Doxo+SAHA-treated group compared to the Doxo group, whereas wall thicknesses were decreased after doxorubicin application but normalized with SAHA co-treatment (**Figures 3H-K**). On histology cardiac fibrosis was increased in Doxo-treated mice but blunted by SAHA co-treatment. **(Figure 3L-M).** Transcriptional analysis of heart tissue 10 weeks after the last doxorubicin application revealed a distinct pattern of differentially enriched genes, which was not observed with SAHA co-treatment. Notably, *Myh7*, which we identified as a Topo IIb/MEF2 top target was among the significant upregulated genes after doxorubicin **(Figure 3N).**

**Figure 3.**
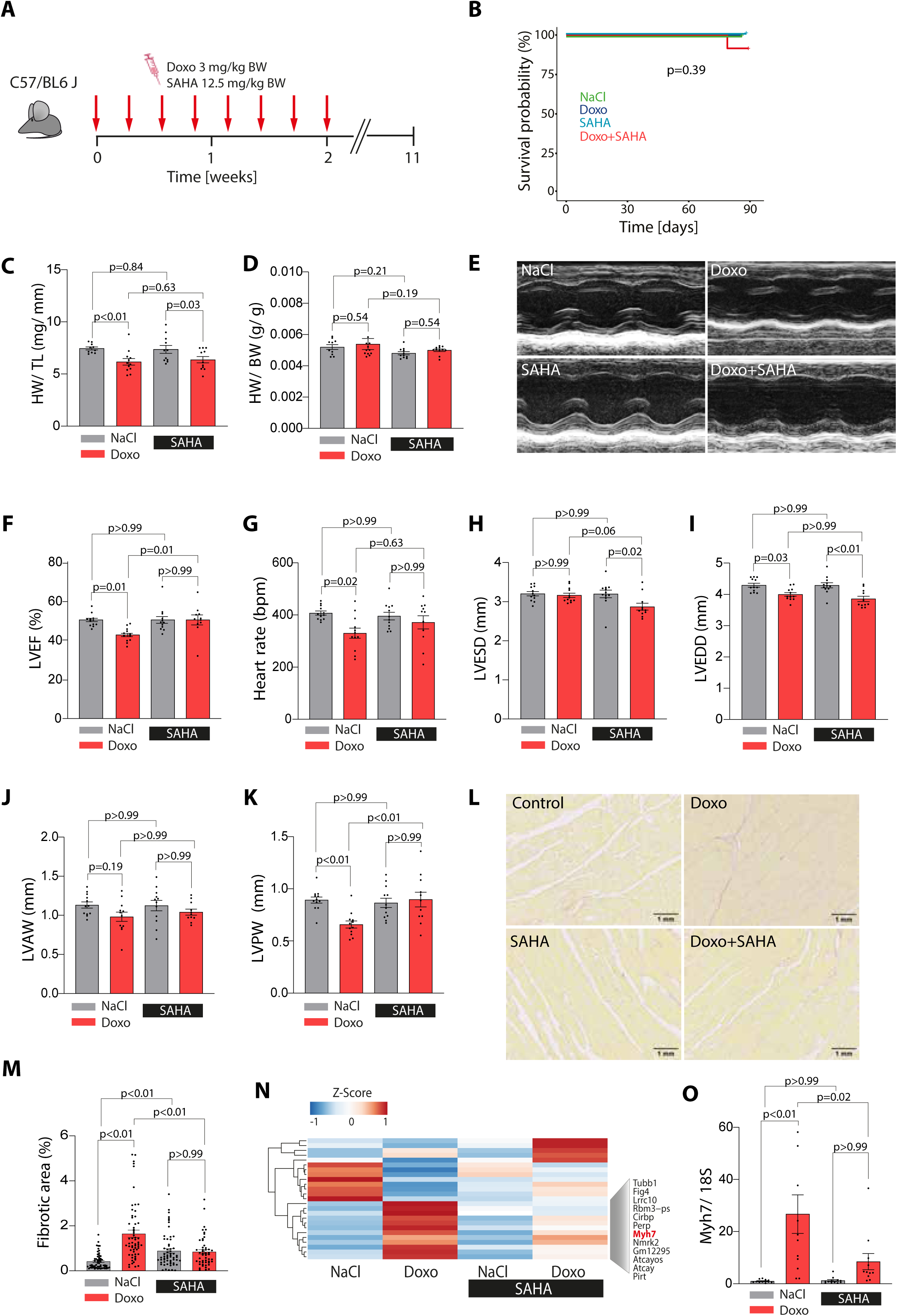
(**A**) Graphical illustration of the in-vivo model of doxorubicin (Doxo) and SAHA applications in C56/BL6J mice as indicated **(B)** Kaplan Meier curve to show survival probability (%) over time. P-value calculated via log-rank test. **(C)** Heart weight (HW) normalized to tibia length (TL) and **(D)** to bodyweight (BW) **(E)** Exemplary images of echocardiography (M-Mode, short axis, anaesthesia with 2% isoflurane). **(F)** Left ventricular ejection fraction (LVEF) was reduced in Doxo-treated mice and not in SAHA co-treated animals (Control 50.32±3.11% vs. Doxo 42.96±3.44%%; p<0.05, n=12; vs. Doxo+SAHA 50.61±7.4%, p>0.05, n=11, Anova with Bonferroni posttest) **(G)** Heart rate in beats per minute (bpm) **(H)** Left ventricular end systolic diameter (LVESD) in mm **(I)** Left ventricular end diastolic diameter (LVEDD) in mm **(J)** Left ventricular anterior wall (LVAW) in mm **(K)** Left ventricular posterior wall (LVPW) in mm. **(L)** Exemplary images of fibrosis with Picro Sirius Red staining. **(M)** Quantification of fibrosis area (%). Anova with Bonferroni posttest **(N)** Differentially enriched genes (DEGs, FDR < 0.05) between Doxo and NaCl treatment which are further normalized after SAHA treatment. Single gene names are shown. *Myh7* is marked in red. Genes with log2 fc >1 or < –1 are shown. Heatmap represents the Z-score of the mean value per group (n=3 animals/group). **(O)** *Myh7* expression measured via qPCR, normalized to 18S expression (NaCl: n=12; Doxo: n=11; SAHA: n=12, Doxo+SAHA: n=11). Anova with Bonferroni posttest.

#### The selective class II HDAC inhibitor TMP195 aggravates doxorubicin-induced cardiotoxicity

To further evaluate the cardioprotective potential of class II HDACs, SAHA was replaced with the selective class II histone deacetylase (HDAC) inhibitor TMP195 (15 mg/kg), which specifically inhibits HDAC4, HDAC5, HDAC7 and HDAC9. The experimental protocol was slightly modified in duration and treatment dosages (**Figure 4A**), since animals treated with Doxo + TMP195 showed higher mortality within the first two weeks of TMP195 application compared to the TMP195 only group (**Figure 4B**). The decrease in heart weight normalized to tibia length after doxorubicin treatment was slightly deteriorated with TMP195 co-treatment (**Figure 4C**), with a decrease in bodyweight in the doxorubicin-treated groups (**Figure 4D, Suppl. Figure 5**). LVEF was significantly reduced by doxorubicin and declined further in the Doxo + TMP195 group **(Figure 4E, G)**, while heart rate was unchanged **(Figure 4F)**. LVEDD and wall thickness were also decreased in the Doxo + TMP195 groups relative to NaCl controls **(Figure 4H–K)**. TMP195 did not alleviate doxorubicin-induced cardiac fibrosis **(Figure 4L–M)**, and TMP195 alone had no significant effects on fibrosis or cardiac function compared with the NaCl group. Additionally, *Myh7* expression, used as a readout of MEF2 activation, was significantly increased in both the Doxo and Doxo + TMP195 groups **(Figure 4N)**.

**Figure 4.**
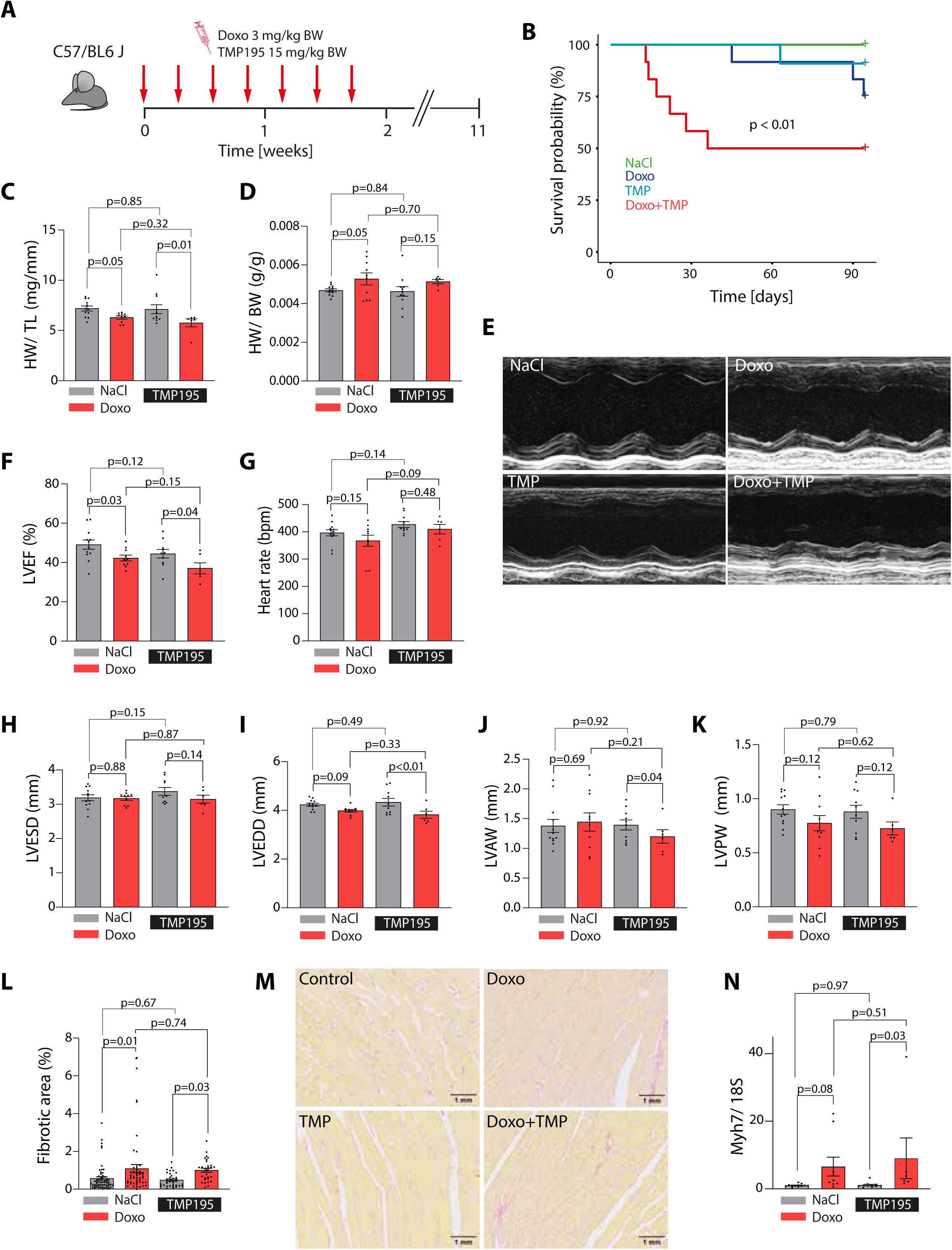
(**A**) Graphical illustration of the in-vivo model of Doxorubicin (Doxo) and TMP195 (class II HDAC inhibitor) applications in C57/BL6J mice **(B)** Kaplan Meier survival analysis shows a significant decrease in survival probability in Doxo/TMP195 co-treatment (log-rank test) **(C)** Heart weight (HW) of the animals that survived normalized to tibia length (TL) and **(D)** to body weight (BW, NaCl: n=12, Doxo: n=10, TMP195: n=11; Doxo+TMP195: n=6)) **(E)** Representative images of echocardiography (M-Mode, short axis, anaesthesia with 2% isofluran). **(F)** Left ventricular ejection fraction (LVEF) shows a reduction in Doxo and Doxo+TMP group (Anova with Bonferroni posttest). **(G)** Heart rate in beats per minute (bpm) **(H)** Left ventricular end systolic diameter (LVESD) in mm **(I)** Left ventricular end diastolic diameter (LVEDD) in mm **(J)** Left ventricular anterior wall (LVAW) in mm **(K)** Left ventricular posterior wall (LVPW) in mm. **(L)** Quantification of fibrotic area reveals increase in Doxo-and Doxo+TMP195-treated mice. Anova with Bonferroni posttest. **(M)** Exemplary images of fibrosis Picro Sirius Red staining. **(N)** *Myh7* expression measured via qPCR, normalized to 18S expression (NaCl: n=12; Doxo: n=9; TMP195: n=11, Doxo+TMP195: n=6). Anova with Bonferroni posttest.

#### Protective effects of SAHA depend on presence of HDAC4

Based on the divergent effects observed with pharmacological class II HDAC inhibition, we next tested whether the cardioprotective effects of SAHA depend on the presence of HDAC4 using a genetic approach. We used a conditional cardiomyocyte-specific knockout mouse model for HDAC4 (HDAC4-cKO)^21^. Efficient deletion of HDAC4 was confirmed by western blotting (**Suppl. Figure 6)**. In the following the animals were treated with doxorubicin (21 mg/kg in 2 weeks) and SAHA (87.5 mg/kg in 2 weeks; **Figure 5A**). HDAC4-cKO mice treated with doxorubicin exhibited a significant reduction in survival which could not be rescued by co-treatment with SAHA (**Figure 5B**). Due to the increased mortality in this genotype, the observation period was shortened from 9 weeks to 5 weeks. During the period, HDAC4-cKO mice showed pronounced weight loss (**Suppl. Figure 7A**) whereas no relevant changes in heart weight were observed (**Figure 5C, Supp. Figure 7B).** In contrast to WT animals, SAHA failed to improve LVEF in doxorubicin-treated HDAC4-cKO after 5 weeks (**Figure 5D-E**). Heart rate tended to be lower in treated HDAC4-cKO animals (**Suppl. Figure 7C**). Moreover, LVEDD and LVESD were significantly higher in HDAC4-cKO following SAHA and doxorubicin co-treatment, even compared to doxorubicin treatment alone (**Suppl. Figure 7D, E**). Wall thicknesses were not altered significantly (**Suppl. Figure 7F, G**). Notably, doxorubicin induced fibrosis was further exacerbated in HDAC4-cKO upon SAHA co-treatment (**Figure 5F-G**). To assess the transcriptional consequences underlying these phenotypes, we performed RNA-seq analysis. PCAs are shown in **Suppl. Figure 7H-J** (Two outliers were excluded from the analysis.). Differential expression analysis revealed a distinct gene expression signature (**Suppl. Figure 7 K**) that was induced after doxorubicin and normalized by SAHA in control littermates, but remain highly activated in HDAC4-cKO despite SAHA treatment (**Figure 5H**). Transcriptional networks derived from GO-terms of this persistently activated gene cluster (marked with a red square) revealed strong enrichment of fibrosis-associated pathways, including ‘response to mechanical stimulus’, ‘extracellular matrix organization’ or ‘cellular response to transforming growth factor beta’. Consistent with our previous data, *Myh7* was found again within this gene cluster, next to e.g., collagens (*Col8a1, Col1a2*) and remodelling associated transcripts, such as *Mmp23, Adamts2/8* or *Ankrd1* (**Figure 5I, Suppl. Figure 7L**). Looking specifically again at *Myh7*, the course of its regulation resembles the regulation of cardiac fibrosis and showed additional upregulation HDAC4-cKO animals receiving combined doxorubicin and SAHA treatment (**Figure 5J**).

**Figure 5.**
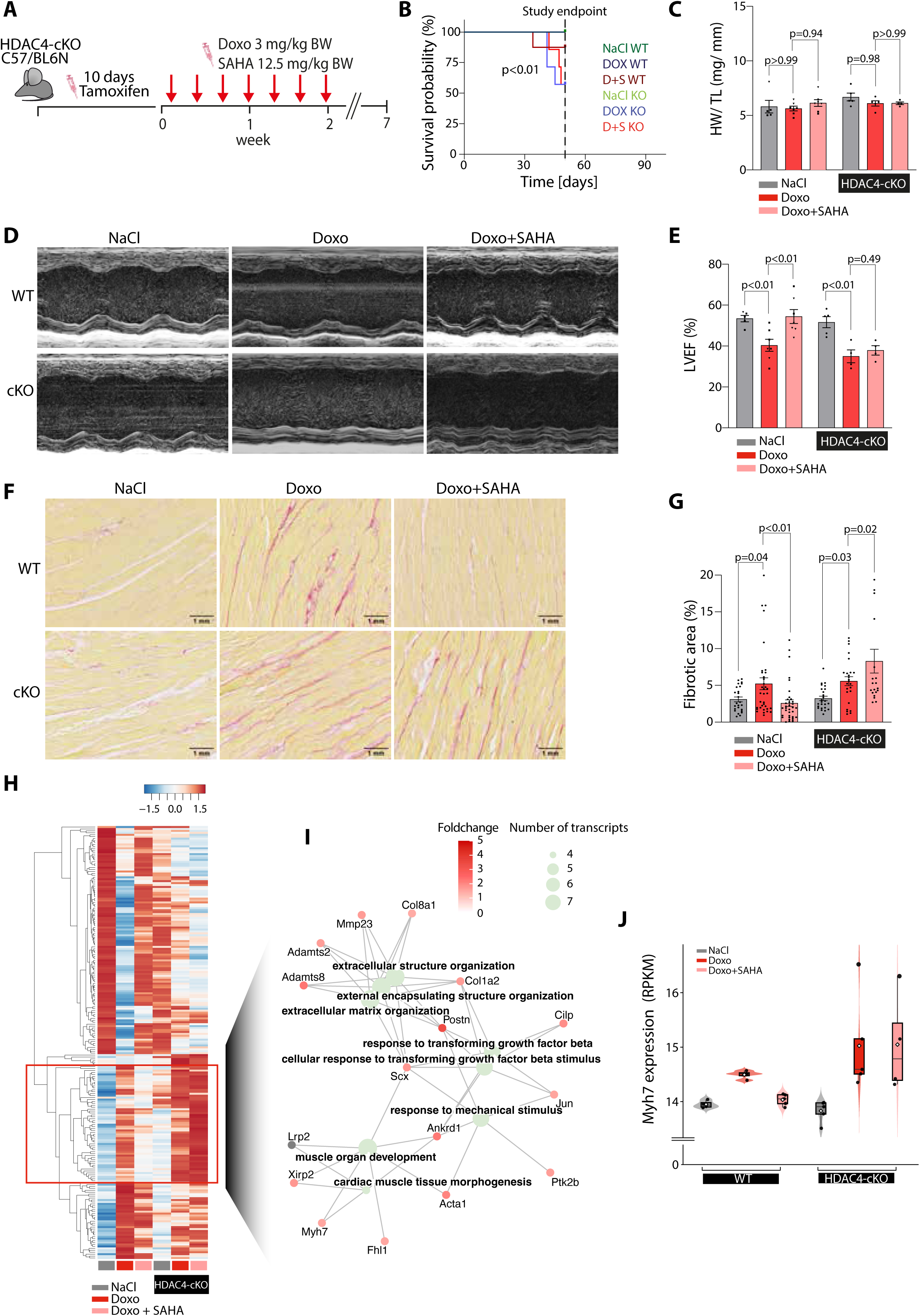
(**A**) Graphical illustration of the in-vivo model in HDAC4-cKO mice and wildtype control littermates (WT) that were treated with doxorubicin (Doxo) and the co-treatment Doxo/SAHA. **(B)** Kaplan Meier survival analysis shows a significant decrease in the survival probability in HDAC4-cKO animals that were treated with Doxo ± SAHA (log-rank test) **(C)** Heart weight (HW) of the animals that survived normalized to tibia length (TL; WT NaCl: n=5; WT Doxo: n=7; WT Doxo+SAHA: n=7; HDAC4-cKO NaCl: n=5; HDAC4-cKO Doxo: n=5; HDAC4-cKO Doxo+SAHA: n=4; in WT squares: fl^−^/fl^−^ cre+, triangles: fl^+^/fl^+^ cre-) **(D)** Representative images of echocardiography (M-Mode, short axis, anaesthesia with 2% isoflurane) **(E)** Left ventricular ejection fraction (LVEF) shows no cardioprotective effect of SAHA in HDAC4-cKO animals (Anova with Bonferroni posttest). **(F)** Exemplary images of fibrosis staining with Picro Sirius Red **(G)** Quantification of the fibrotic area. Anova with Bonferroni posttest. **(I)** Heatmap of differentially expressed genes (DEG) between NaCl and Doxo treatment in WT animals (FDR <0.05). The gene cluster in the red box highlights genes induced by Doxo that are attenuated by SAHA in WT but not in HDAC4-cKO animals. **(I)** Gene Ontology (GO) network of the genes in this cluster. Node colour reflects the fold-change (Doxo vs. NaCl) and node size indicates the number of transcripts contributing to each GO term. **(J)** Violin plot shows relative *Myh7* expression (RPKM) in all groups.

## Discussion

Cardiotoxic side effects remain a major limitation to cancer therapies creating an urgent need for less harmful drugs^29^ or effective cardioprotective co-therapies. Although oxidative stress and Topoisomerase IIb – dependent DNA damage have been implicated, the mechanisms by which these events are translated into sustained maladaptive gene programs driving cardiac remodeling have remained poorly understood. In this study, we identify a previously unrecognized epigenetic–transcriptional axis that links doxorubicin-induced DNA damage to MEF2-dependent pathological remodeling in cardiomyocytes. In this regard, HDACis, as a relatively new class of anti-cancer drugs, have been shown to exert cardioprotective effects in several preclinical cardiac stress models^5,11,30–32^. However, the specific mechanism how HDACis are protecting from pathological cardiac remodeling and cardiac dysfunction remained largely unresolved.

Our findings suggest that doxorubicin-poised Topo IIb DNA binding preferentially affects MEF2-regeulated genomic regions, indicating that anthracyclines not merely cause random genotoxic stress but instead target transcriptional relevant regions, critical for cardiomyocyte stress response. This directly connects doxorubicin-associated activation of maladaptive gene programs (e.g., hypertrophy, fibrosis) and defined MEF2 as a central integrator DNA damage.

We identified the acetylation of 14-3-3 as critical step to regulate nuclear retention of class II HDACs and maintained MEF2 repression. This highlights non-histone protein acetylation as an essential and previously underappreciated mechanisms in cardiac stress signaling.

The functional importance of this pathway is underscored by the complete loss of cardioprotective effects of SAHA in cardiomyocyte-specific HDAC4-deficient mice. In this model, SAHA fails to suppress pathological remodeling and instead exacerbates fibrotic response, demonstrating that HDAC4 is and indispensable mediator of SAHA-dependent cardioprotection. Increased mortalty and absence of cardioprotective effects in the HDAC4-specific inhibitory drug askes for a unresolved non-canonical and potentially cytosolic function of HDAC4. The increased mortality was not explained by heart failure and specific causes of death were not further evaluated. Due to the combinational use of pharmacological inhibitors, any organs system may be involved.

The inhibition of HDACs by pan-inhibitors has been shown earlier to have cardioprotective effects in other settings.^5,6^ Here, we extend these finding by demonstrating a cardioprotective effect of pan-HDACi in a model of anthracycline-induced cardiotoxicity in vivo. Given its established antiproliferative activity, SAHA represents an attractive candidate for repurposing as a cardioprotective co-therapy in oncological treatment strategies. Importantly, MEF2 activity was in inhibited in NRVMs at a SAHA concentration of 3µM, levels comparable to serum concentrations observed in patients following oral SAHA administration^33,9^.

However, we cannot exclude the possibility that other targets, inhibited by SAHA^28^ may also contribute to the cardioprotective effects in cardiomyocytes. HDAC6 was shown to be maladaptive in the heart and mediate muscle wasting in response to Angiotensin II in a mouse model. A selective inhibitor of HDAC6, tubastatin A, could show similar protective effects compared to HDAC6 knockout animals.^14^ These results support our finding of MEF2 inhibition in HDAC6 knockdown cells. Following our model, the effect of HDAC6 can be explained via acetylation of 14-3-3 and the following enhancement of the cardioprotective effects of HDAC4 and 5.

Considering the impact of pan-HDACi or selective inhibitors on the heart, the opposing or synergistic roles of several HDACs and their interaction among each other have to be taken into account in the treatment of heart failure. Following our findings, the positive effect of SAHA is presumably due to its mode of action to inhibit the deacetylase subunit, which is most relevant in class I HDACs and inhibits less class II HDACs in its protein binding ability.^16^

In regard to therapies of malignant diseases including different HDACis, the cardiac influence of the inhibited HDACs should be considered.

Clinically, these findings have important implications. They provide a mechanistic rationale for the use of pan-HDAC inhibitors as cardioprotective co-therapies during anthracycline treatment, while cautioning against selective inhibition of class II HDACs^34^.Our study identifies HDAC4-14-3-3 acetylation as a critical node in chemotherapy-induced cardiac remodeling and a promising therapeutic approach regarding the development of more specific cardioprotective strategies.

